# Antibiotic Susceptibility Profile of *Campylobacter Spp* from Poultry

**DOI:** 10.1101/2021.04.09.439140

**Authors:** Khalda A. Khalifa, Elbrissi Atif, Igbal S. Adel Rahim, Amgad M. Abdel Rahman, Salwa A. Ahmed

## Abstract

Campylobacteriosis is found among the four important worldwide food borne pathogens. In intensive poultry rearing systems in Sudan the use of oral antibiotics is essential to preserve health. Accordingly, there is a high threat for the thermophilic *Campylobacter spp* inhabitant in the intestinal tract of food animals to develop resistance to commonly used antibiotics. Contamination of broiler meat with pathogenic strains of resistant *Campylobacter* could, therefore, result in a form of campylobacteriosis in humans that is difficult to treat. To the best of our knowledge, there are no data available relating antibiotic resistant against these bacteria in poultry in Sudan, hence the aim of this study was to determine the antimicrobial susceptibility profile of thermophilic *Campylobacter* spp. isolated from broiler in Khartoum by disk diffusion. Sensitivity of fourteen isolates against seven antibiotics namely Neomycin, Nitrofurantoin, Nalidixic acid, Gentamycin, Streptomycin, Tetracycline and Erythromycin were studied. The result showed that Nitrofurantoin was found as highly sensitive (64.3%). It was also observed that Erythromycin, Gentamycin, Streptomycin and Neomycin verified the following resistant against isolates recovered 92.9%, 71.4%, 71.4%, 64.3% respectively. Multidrug resistant against five, four, three and two antibiotics was also reported. It was concluded that poultry meat in Khartoum could complicate the antimicrobial therapy in human as multidrug resistant of campylobacter was detected.

## Introduction

Poultry industry in Sudan witnessed considerable development only in the last 10 years, with production increasing from 5 million broilers in 2006 to close to 90 million in 2017. While several factors contributed to this increase, the two most important were the government decision to stop imports of frozen poultry in 2006 and the increase in red meat prices. Other factors that contribute to increase of poultry meat consumption are, urbanization, change in food habits, rising income and population growth [1].

Increasing meat consumption worldwide raises a lot of concerns and challenges to meat hygiene and safety [2]. One of the major concerns is contamination of meat by products by campylobacter particularly poultry meat. *Campylobacter spp* reside in the gut of domesticated animals and birds as part of the intestinal microflora [3]. The Liver and heart of chickens’ carcass (known as giblets) are separated from the slaughtered bird during processing and often packaged with them. Giblets may also be purchased separately as livers, hearts, or a combination.

*Campylobacter* spp. is Gram negative, microaerophilic, curved or spiral rods in the family Campylobacteriaceae [4].

Campylobacter is considered as one of the most common pathogen-related causes in diarrheal illness globally and has been recognized as a significant factor of human diseases over three decades [5]. Most farmers in Sudan chose antibiotics depending on availability and cost and/or what the local sales person recommended rather than consulting a veterinarian. Knowledge of resistance to antibiotics was generally low. This study was conducted to find out the antibiotic susceptibility profile and resistant as no data was present.

## Materials and Methods

### Source of samples

Samples (livers, heart and intestines) of slaughtered broiler chickens were collected from fourteen slaughterhouses located in 5 different localities in Khartoum State namely (Gebel awlia1, bahri Alkadroo, Gebel awlia2, bahri Alsagai and Sharq alneel) (Table 1), during the year 2020. Three visits were made to each slaughterhouse with exception of one where visited twice. Samples of livers, hearts and intestines organs were collected during slaughtering, pooled, packed in sterile bags and then transferred on ice bags to the lab.

**Table (1).** Antibiotic susceptibility of campylobacters isolated from different localities. Total sensitivity tests according to location was illustrated in table (1). Percentage of resistance ranges between 38.1 in SharqAlneel and 71.4 in Bahri Alsagai while total resistance was found to be 54.1. Sensitivity to the mentioned antibiotics was high (42.9%) in SharqAlneel. Moderate responses revealed by the antibiotics were nearly the same (table 1).

| *Location* | *No of isolates* | *No. sensitive (%)* | *No. moderate (%)* | *No. resistant (%)* | *Total sensitivity test (%)* |
| --- | --- | --- | --- | --- | --- |
| *Gebel Awlia 1* | 2 | 4/14(28.6) | 3/14 (21.4) | 7/14 (50) | 14/98 (14.4) |
| *Bahri Alkadaro* | 3 | 5/21(23.8) | 6/21(28.6) | 10/21(47.6) | 21/98 (21.4) |
| *Gebel Awlia 2* | 3 | 3/21(14.3) | 5/21(23.8) | 13/21(61.9) | 21/98(21.4) |
| *Bahri Alsagai* | 3 | 1/21(4.8) | 5/21(23.8) | 15/21(71.4) | 21/98(21.4) |
| *SharqAlneel* | 3 | 9/21(42.9) | 4/21(19) | 8/21(38.1) | 21/98(21.4) |
| *Total* | 14 | 22/98(22.4) | 23/98(23.5) | 53/98(54.1) | 98 |

### Bacteriological examination

Collected pooled samples of each visit were incubated in Cary Blair medium overnight at 42 °C, then streaked onto Skirrow’s agar plates and charcoal-based media (CampyGen, Oxoid, CN35A) supplemented with antibiotic supplement mixture (vancomycin, polymyxin B and trimethoprim) were added to the media for primary isolation and incubated at 42°C and 37°C for 48-72 h under micoraerophilic conditions in an anaerobic jar (CampyGen, CN0225, Oxoid Ltd).Representatives of typical and well-isolated colonies were sub-cultured onto nutrient agar plates for purification which was checked by Gram staining. Heavy suspension of 24-h growth for each pure isolate was prepared in 10 ml of 20% glycerol-peptone medium, mixed well and suspended in 2.5 ml volume into four vials. All suspended colonies were identified by microscopic examination of morphology and motility, oxidase test, catalase test and standard biochemical methods as described previously [6].

### Antibiotic Susceptibility Test

Resistance of *Campylobacter* isolates to certain antibiotics was determined by Kirby-Bauer disk diffusion method on Mueller-Hinton agar (Liofilchem, Italy) supplemented with 5% sheep blood following CLSI 2016 guidelines. Seven antibiotics tested and their corresponding concentrations were Neomycin (30 mg), Nitrofurantoin (300 mg), Nalidixic acid (30 mg), Gentamycin (10 mg), Streptomycin (10 mg), Tetracycline (30 mg), and Erythromycin (25 mg). For the disk diffusion method, sterile cotton-tipped swabs were used to transfer the inoculum onto Mueller-Hinton plates to produce a confluent lawn of bacterial growth. After the inoculum on the plates was dried, the above mentioned antibiotic disks were distributed over the inoculated plates. The inhibition zones were recorded and interpreted following CLSI breakpoints.

## Results

### Identification of isolated bacteria

All collected samples (livers, hearts, and intestines) revealed *Campylobacter spp* by cultural methods. Colonies of *Campylobacter spp* on Skrirrow selective medium appeared as pink while light grey colonies on charcoal-based medium. Microscopic examinations revealed Gram-negative small spiral curved bacilli with comma-shaped (S) or gull wing-shaped cells that showed high motility and corkscrew like, all isolates were oxidase and catalase positive.14 *campylobacter spp* isolates were recovered from the five mentioned slaughterhouses as in table 1.

### Antibiotic susceptibility Test

Sensitivity of each isolates against seven antibiotics namely Neomycin, Nitrofurantoin, Nalidixic acid, Gentamycin, Streptomycin, Tetracycline and Erythromycin ranged between (0-71.4%). Moderate responses ranged between (0-57.1%) while resistant varied between (14.3%-85.7%) as in figure 1.

**Figure:(1).**
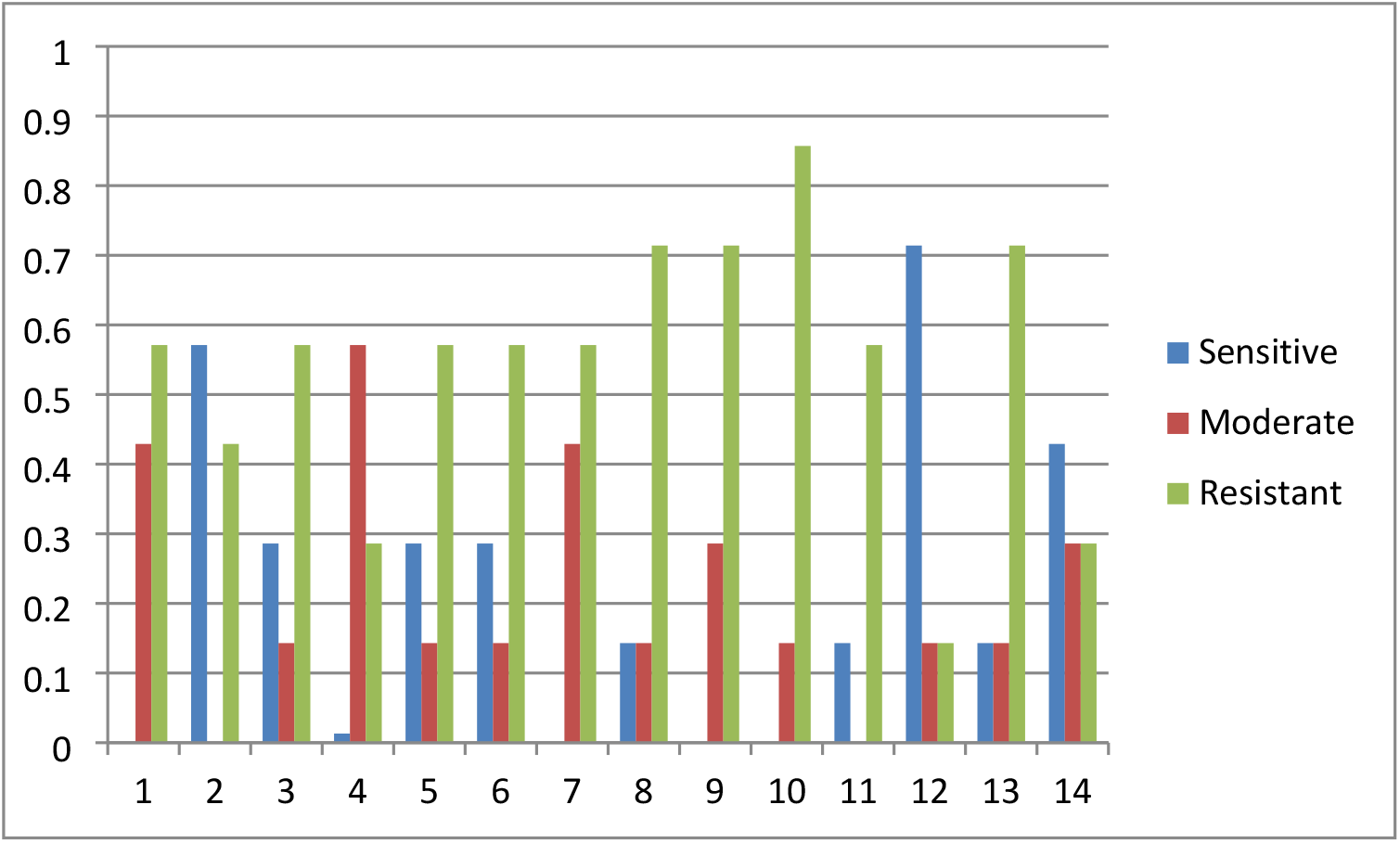
Antibiotic sensitivity profile of Campylobacter Isolates against the seven antibiotics. Nitrofurantoin was found as highly sensitive (64.3%) among antibiotics examined and tetracycline (7.1%) as less sensitive against campylobacter detected. Tetracycline revealed 64.3% high moderate whereas Erythromycin recorded 7.1%. It was also noticed that Erythromycin, Gentamycin, Streptomycin and Neomycin verified the following resistant against isolates recovered 92.9%, 71.4%, 71.4%, 64.3% respectively as in figure 2.

**Figure (2).**
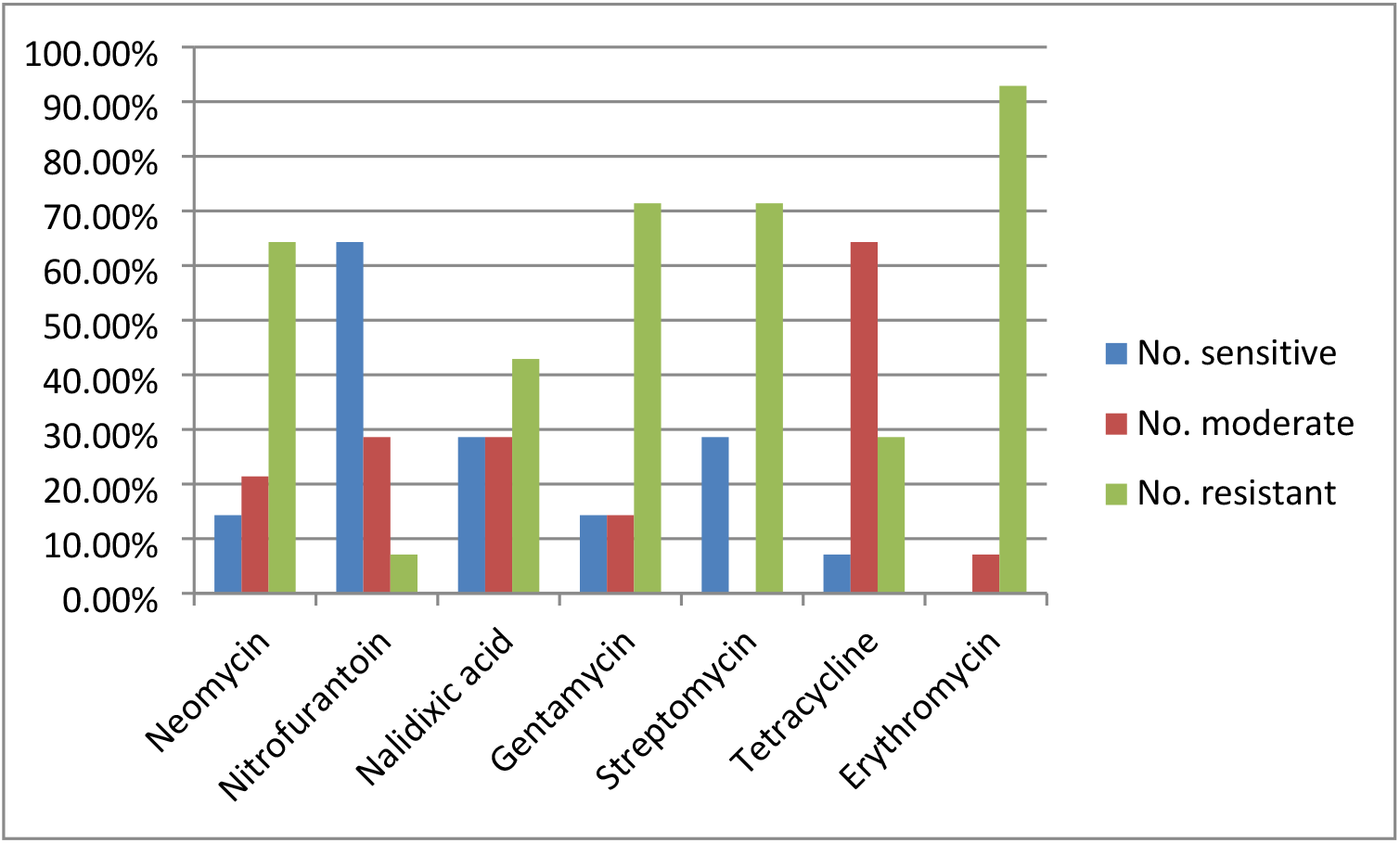
Sensitivity patterns of the seven antibiotics. In the current study multidrug resistant (MDR) against five, four, three and two antibiotics were reported.10 isolates were found resistant to Gentamycin; Streptomycin and Erythromycin.10 isolates were resistant against Streptomycin and Erythromycin. 9 isolates resistant against Neomycin and Erythromycin, the details in figure 3.

**Figure (3).**
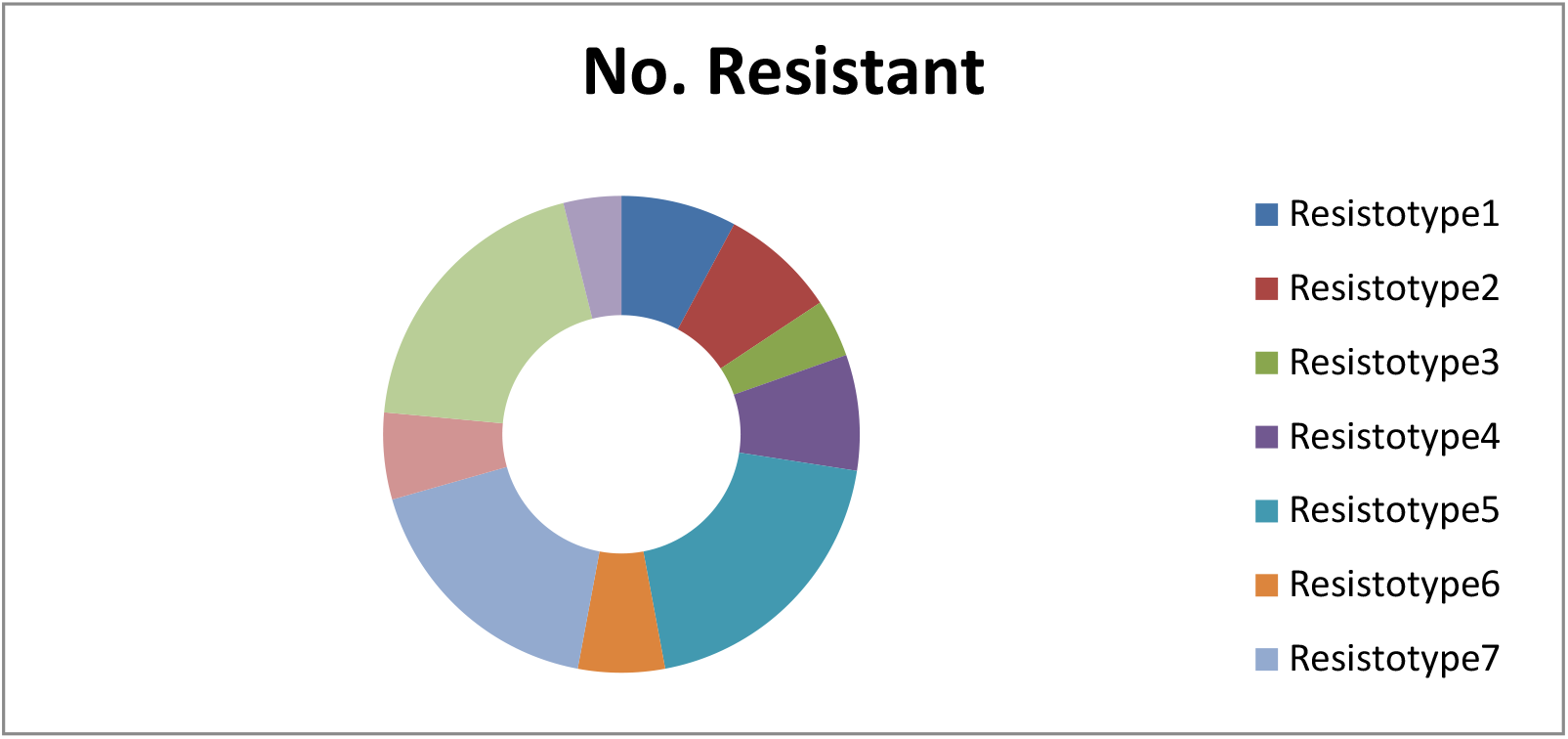
Multi-drug resistant of Campylobacter isolates. Resistotype 1: Neo Nalid Gent Strept Erythro, Resistotype 2: Neo Gent Strept Erythro, Resistotype 3: Nalid Gent Strept Erythro, Resistotype 4: Neo Gent Strepto, Resistotype 5: Gent Strepto Erythro, Resistotype 6: Nalid ErythroResistotype 7: Neo Erythro, Resistotype 8: Tetra Erythro, Resistotype 9: Strepto Erythro, Resistotype 10: Strepto Tetra.

## Discussion

Poultry meat is considered the most important source of human *Campylobacteriosis* and the role of poultry as a reservoir for transmission to humans has been recognized. Around 20%–30% of human infections are linked to the manipulation, preparation and consumption of broiler meat, while 50%– 80% may be attributed to the chicken reservoir as a whole [7].

*Campylobacter* spp. represents human health risks when poultry is inadequately cooked or is cross-contaminated after cooking [8]. Cross-contamination can occur at each phase in the process of bringing the product to the consumer, beginning at the farm level and continuing through processing [8]. Slaughtering and processing steps such as scalding, picking, and chilling may be a source of cross-contamination [9]. Additionally, meat can become contaminated with pathogens from intestinal contents, skin, or feathers [9].

High degree of sensitivity to Erythromycin, resistant 72.7% to tetracycline from 33 campylobacter isolates from Arabian Gulf was reported [10]. They also reported MDR in 6 poultry isolates in comparison with 92.9% Erythromycin resistant and 10 isolates as maximum MDR in the current study.

Resistant to Erythromycin 7%, Tetracycline 71%, Streptomycin 21%, Gentamicin 7% and Nalidixic acid 71% was reported in Vitnam [11] compared to 92.9%,28.6%,71.4%,71.4% and 42.9% respectively in this study.

Therapeutic antimicrobials of choice in human patients suffering from life threatening campylobacteriosis are initially the macrolides and thereafter the fluoroquinolones and Gentamicin [12]. Resistance to these 2 classes of antibiotics in zoonotic *Campylobacter* species can increase hospitalization rates and the cost of therapy, and decrease the survival rate of patients [12]. A study in South Africa revealed there was no resistance in the *Campylobacter* spp. isolated from birds in Gauteng to Gentamicin, neomycin that disagreed with what was found here [13].

In the current study, multidrug resistant (defined as resistance to four or more different classes of antibiotics) is found in 10 strains as max, in contrast to 4 strains detected by [13] despite no multiple resistance has been also reported in a number of countries—four European Union countries, Northern Ireland and Sweden [14,15,16] The reasons of these results may be due to the fact that urbanization and increase consumption of poultry meat besides time interval of earlier studies. Another study reported (14.8%-19.2%) Tetracycline, Erythromycin 11.1% and no resistant showed by Nalidixic acid from poultry in the South-East Queensland region [17]. Higher levels of tetracycline resistance have been reported from four European Union countries (35.4%) and the USA (43%) [14,18]. In Sweden, where tetracycline has not been added to chicken feed since 1986 [16]. The level of tetracycline-resistant Campylobacter has been reported to be 1% [16,17]. A study of 62 broiler strains in Brazil revealed resistant to Neomycin (50%), Tetracycline (43%) whereas a lower percentage of strains was resistant to Erythromycin (10%), all stains were sensitive to Gentamycin [19]. highest rate of resistance (75 %) Nalidixic acid, while Gentamycin was the most active (96%) antibiotic against studied isolates followed erythromycin (95%) and streptomycin (94%) in chicken and beef meat in Iran [20]. Unlike this study, total resistance of campylobacter isolates to Nalidixic acid and total sensitive to erythromycin in Indian poultry [21].

As multidrug resistant was detected here; constant awareness for *Campylobacter* spp. of public health significance should be maintained through the use of surveillance and the rapid reporting of trends. Previous study [7] reported 81.45%, 3.76%, 75.6%,22.96% in comparison to42.9%,71.4%,28.6% and 92.9% of Nalidixic, Gentamycin, Tetracycline and Erythromycin in the current study. The regional differences in the prevalence of AMR among the Campylobacter isolates in the present study could be the result of the manner in which antimicrobials are used in veterinary medicine. There are differences in production and veterinary practices between regions with regard to drug prescription, sales, or marketing since those matters are governed by provincial laws [22] and resistance profiles for chickens reflected the drugs administered on the farm [23]. Environmental factors that are specific to certain regions could also be a source of resistant Campylobacter. In conclusions, the high rate of contamination in chicken meat in Khartoum alarms a significant public health concern. Most of the isolates were resistant; therefore, there is a possible risk of human infection with *Campylobacter* spp. via consumption of these products.

